# CellBoost: A pipeline for machine assisted annotation in neuroanatomy

**DOI:** 10.1101/2023.09.13.557658

**Authors:** Kui Qian, Beth Friedman, Jun Takatoh, Fan Wang, David Kleinfeld, Yoav Freund

## Abstract

One of the important yet labor intensive tasks in neuroanatomy is the identification of select populations of cells. Current high-throughput techniques enable marking cells with histochemical fluorescent molecules as well as through the genetic expression of fluorescent proteins. Modern scanning microscopes allow high resolution multi-channel imaging of the mechanically or optically sectioned brain with thousands of marked cells per square millimeter. Manual identification of all marked cells is prohibitively time consuming. At the same time, simple segmentation algorithms suffer from high error rates and sensitivity to variation in fluorescent intensity and spatial distribution. We present a methodology that combines human judgement and machine learning that serves to significantly reduce the labor of the anatomist while improving the consistency of the annotation. As a demonstration, we analyzed murine brains with marked premotor neurons in the brainstem. We compared the error rate of our method to the disagreement rate among human anatomists. This comparison shows that our method can reduce the time to annotate by as much as ten-fold without significantly increasing the rate of errors. We show that our method achieves significant reduction in labor while achieving an accuracy that is similar to the level of agreement between different anatomists.

## 1 Main

We present an adaptive system designed to assist neuroanatomists with the task of annotating cells labeled with a molecular label. The standard, fully manual approach, referred to here as “unaided”, requires anatomists to detect as many marked cells as possible in each section cut from the brain. In our approach, a computer detector identifies confident detections versus unconfident detections of potentially labeled cells across each section. The anatomist then performs two tasks. The first is quality control (QC) on the set of confident detections. Here the anatomist examines, and either confirms or disputes, the computer’s identification for a small fraction of this set of cells. The second task is to examine all unconfident detections of cells; this corresponds to labeled cells that are difficult to categorize. In this way, the major time of the anatomist is spent attending to the typically small population of cells whose labeling is ambiguous. The reduction in anatomist labor therefore depends on the ratio between the automated confident and unconfident detections.

### 1.1 Significance for Neuroscience

How do we study the computational pathways of brains? Brains are composed of circuits at multiple scales of organization. These range from the one to ten micrometer-scale of connections between individual neurons, i.e., the microscopic, to the hundreds of micrometers to millimeters scale of collections of neurons that form computational units, i.e., the mesoscopic. Molecular tagging of neuronal cell types by the expression of genetically encoded reporters and light-level imaging of the cells and tags, using both transmission and fluorescent microscopy, plays an essential role in this process. The resulting raw data files are often extremely large, from 1 to 100 tera-byte per mouse brain. The current means for quantification and spatial mapping of tagged neuronal populations in brain sections require labor- and time-intensive manual annotation by expert neuroanatomists. How can machine learning assist with this process and minimize the amount of manual labor involved, while maintaining accuracy and consistency?

Our approach to neuroanatomy uses mouse brains that are serially sectioned and stained with two markers. The somata of each cell is stained with one fluorophore, a process denoted counterstaining. In addition, specific cell types are marked with a second, different fluorophore based on expression of a designated protein or mRNA. The dual marking approach has the advantage of reducing false detections: stained neurons are identified by combining information from the two separate markers, one as a subset of the other [32]. The images of these two markers for the same cell need not have identical shapes.

### 1.2 Significance to machine learning

The standard goal of a machine learning algorithm is to reach an accuracy as good as, or better than, a human, thereby justifying replacing the human with the learned model. This accuracy is measured according to an agreed upon “ground truth” that associates a “label” with each instance.^1^ Often the labels are subjectively selected by a human, and different humans sometimes assign different labels to the same instance. In such cases the notion of “ground truth” becomes murky and it is unclear how to measure the performance of learned models. This phenomenon, called inter-rater disagreement [19], has been well studied in clinical neuroscience [3] but remains less explored in general neuroscience research. One contribution of this paper is to address the issue of inter-rater disagreement in the context of machine learning for cell detection.

Our approach is to mimic multiple people labeling an instance. We use several learned models, each with a slightly different training set. We test if our models assign different labels to a test instance for which the human assigned labels differ as well. Inter-rater disagreement rates quantify the average level of disagreement. Here we quantify the level of disagreement on each particular instance. We consider the cases of both human and machine labelers. We say that an instance is “sure”, or easily confirmed, if a significant majority of labelers label it the same way. We say that an instance is “unsure”, or difficult to confirm, if significant disagreement exists between labelers. Partitioning instances into “sure” and “unsure” is a central ingredient for a type of learning protocol called active learning [25, 17]. “Sure” and “positive” labeled cells are expected to be have a low false positive rate. To verify that this is actually the case, we take a fixed size sample and have it labeled by multiple human annotators. The key observation here is that only a small sample is needed to verify a low false positive rate.

A main novelty of this paper is that it studies algorithms for cell detection in a dynamic work-flow that combines computers and humans, rather than in isolation, using a fixed training set. The goal is not to generate an autonomous system that can perform cell detection at human accuracy. Instead, the goal is to reduce the amount of human work without degrading accuracy. The details of our approach are given in Section 3.6.

### 1.3 Measures of performance

The methodologies employed to measure the performance of cell segmentation and counting techniques should be both robust and sensitive to the intricacies of cell morphology and the varying conditions under which cells are imaged. Three different metrics and evaluation strategies are typically considered to assess the efficacy and precision of cell segmentation and counting. We adopt one of these.

- Segmentation performance metrics: the Jaccard Index (Intersection over Union), Dice Coefficient (F1 Score), and the Boundary F1 Score. These quantify the spatial overlap and boundary accuracy between the algorithmic predictions and ground truth annotations. Typically, human derived ground truth annotations take enormous labor.
- Counting cells: deviation between the automated counts and the manual counts of labeled cells as performed by human experts. Counting cells in a large brain is time consuming and expertise-dependent. Further, the disagreement between human experts may exceed expectations and need careful reconciliation. Even a matching count between human experts does not imply accuracy of individual cell detection.
- Binary test (adopted here): accuracy rate, false positive rate, and false negative rate of a sample from the automated detections. We reduce human labor by adopting binary annotation for cells instead of using masks. Further, we take a sample from automated detection of cells, and perform QC to provide insight into the accuracy of the automated process.

### 1.4 Comparison to other work

Current popular bio-image analysis tools such as Fiji [24], CellProfiler [20] and so on [37] rely on cell segmentation, which is harder than cell detection. These tools typically use classical segmentation algorithms such as thresholding or watershed. It requires great expertise to select the algorithm that suits the problem and to adjust its parameters. Such manual configurations lead to time-consuming and expertise-dependent processes. Additionally, these classical algorithms may struggle with the heterogeneity of biological samples and technical artifacts, as has been reported in [8, 21, 34]. These limitations hinder the widespread adoption of imaging technologies in biological laboratories [4].

Machine learning-based solutions for cell detection problems exist. Most of these solutions rely on Neural Networks (NNs). Examples include Ilastik [27] and the Trainable Weka Segmentation toolkit of ImageJ [9]. These tools provide plug-in models that take 2D images as input and output binary masks as results. The plug-in models need to be retrained to fit to each experimental situation. The retraining process involves preparing annotations as training data, training models, and configuring algorithms. In particular, preparing annotations is time-consuming, taking humans potentially weeks to draw contours for thousands of cells.

Some efforts have been made to incorporate active learning in this context. Tyson et al. [33] provided a deep learning algorithm for fully automated 3D detection of neuronal somata in mouse whole-brain microscopy images. A traditional image analysis approach, simple thresholding, was used first to detect cell candidates. The list of candidates was then refined by a deep learning step. Harnessing the power of deep learning for object classification rather than cell segmentation at a voxel level speeds up analysis and simplifies the generation of training data. Instead of annotating cell borders in 3D, experts just need to annotate cell candidates by the addition of a single mark. The validation of detected cells was via comparison of cell counts per brain region between the algorithm and the mean of the two expert counts. This required two experts to annotate all marked cell somata throughout the brains. The observed discrepancies between the two experts were noted but not thoroughly examined, ultimately leading to the adoption of their mean assessments for consensus.

In 2018 there was a competition called the Data Science Bowl [4] that used curated training and test data sets. The host team collected and manually segmented a large data set of cell images from a variety of microscopy modalities and fluorescent markers. The competition challenged participants to develop segmentation methods that could be applied to any 2D light microscopy image of stained cell nuclei without manual adjustment. Deep-learning-based models outperformed classical image processing algorithms in terms of accuracy and usability in this competition. The performance of the models was evaluated with the F1 score at different Intersection over Union (IoU) thresholds that served as the primary performance metric. The best-performing solution used an ensemble strategy with eight fully connected convolutional neural network architectures. Models based on the Feature Pyramid Network architecture and the Mask R-CNN (Region-based convolutional neural network) [13] architecture also achieved good performance [4]. Another team followed the approach of the Data Science Bowl team to prepare datasets, and developed a generalist algorithm, called Cellpose [29], for cellular segmentation that can handle diverse cell shapes and image types without requiring model retraining or parameter adjustments. The architecture of Cellpose is described, including the use of simulated diffusion and neural networks to predict spatial gradients and generate masks. We applied this model to our data and compared the results with those found with our method (Figure 1); we contrast the output of Cellpose with both the initial segmentation by CellBoost and the refined, final output of CellBoost.

**Figure 1:**
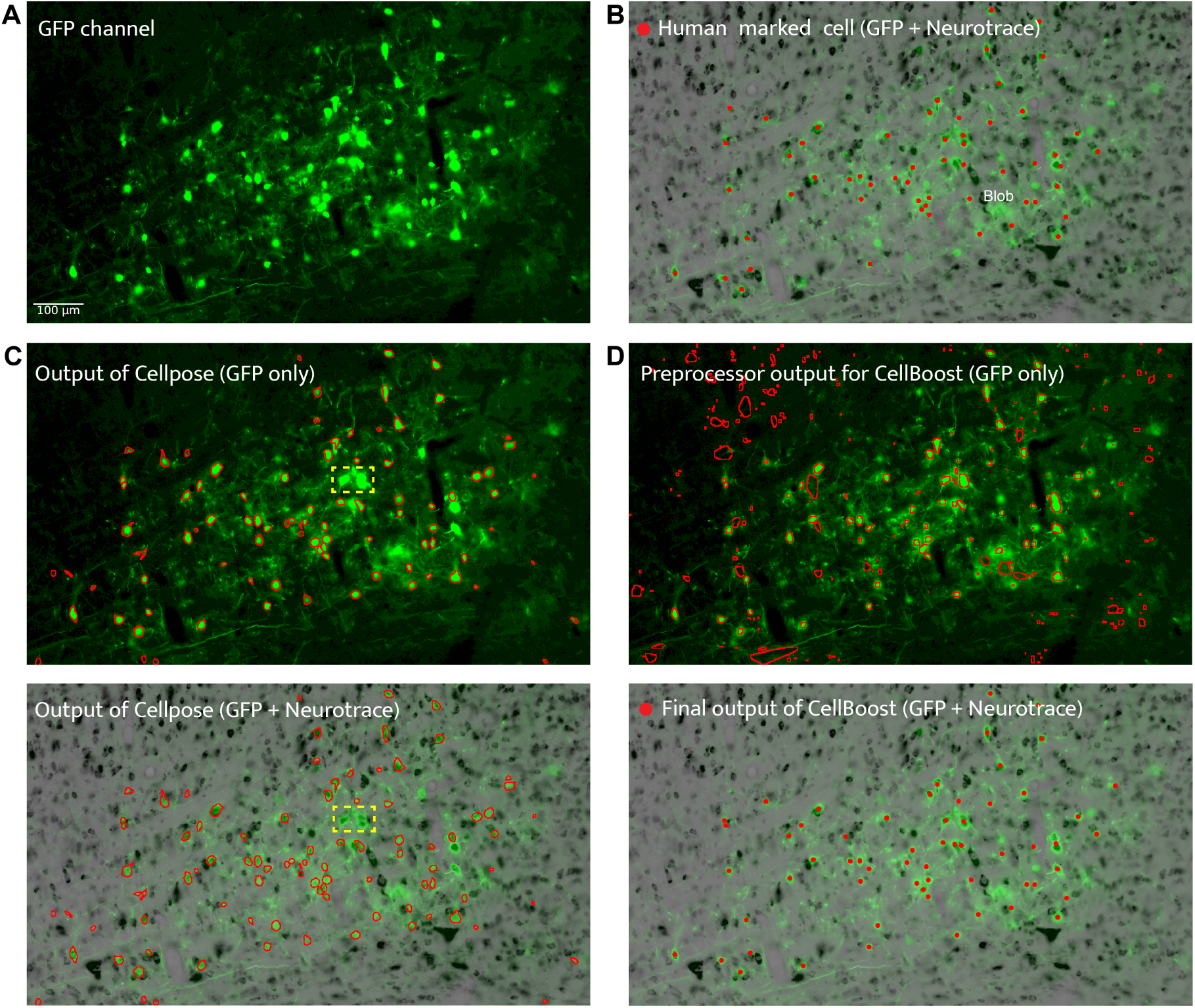
An example of cell detection, as performed by a neuroanatomist and using different segmentation methods. **(A)** A brain section image labeled by GFP. **(B)** A two channel image merging the GFP (green) and the inverted counterstain Neurotrace (gray) images. The red dots correspond to annotations by the anatomist. Note that some of the large green blobs are not judged to be cells. **(C)** Final results of a deep learning segmentation method called Cellpose, using just the GFP channel and both channels, respectively. Cellpose similarly detects most of the labeled cells, but also has high false detection rate, as do most segmentation methods. To remove the false detections reliably, we use a filter created using boosting. The results are also shown over an image with both channels **(D)** Intermediate and final results for CellBoost. The intermediate result is for a simple segmentation method applied only to the GFP channel. Note that all of the human-marked cells are identified, although there are also a large number of false detections. The final result uses both GFP and Neurotrace channels and is close to the human annotation in panel B.

Neural Networks have shown great potential to automate brain section analysis. However, as NN models function as black boxes, the models provide no insight as to how decisions are made, nor do NNs give a rigorous measure of confidence in their predictions. It is thus not possible for a neuroanatomist to understand why the NN made particular decisions. In this paper, we present an alternative approach with models that operate in a way more analogous to that of a neuroanatomist. These models generate outputs that the neuroanatomist can interpret and correct.

An important observation regarding locating labeled cells is that, while a typical brain section will contain some hard to identify features and locations, most of the identifications are relatively easy. This observation drives our methodology, which uses the computer to assist rather than replace the human anatomist. We use confidence rated detectors, which associate a confidence score with each detection. High confidence detections are marked by the computer while low confidence detections are passed on to be be marked by human anatomists.

#### 1.4.1 Human factors and work flow

Accurate identification of neuronal cells is crucial for understanding the functional characteristics of distinct regions within the nervous system. However, selecting appropriate strategies for labeling marked cells can be challenging, especially for non-expert users, as a consequence of the diversity of sample preparation, staining methods, microscopy modalities, and other experimental conditions. In particular, fluorescent labeling comes with undesirable side effects, including photo-bleaching and background signals [36].

We compare the amount of work done by the anatomist when unaided to the amount of work when aided by confidence rated detections. When using an accurate and confident detection system, the work of the anatomist reduces to the following steps:

1. **Performing quality assurance on confident detections:** the anatomist receives a small sampling of the confident detections and verifies that they are correct. The sample size depends on the desired accuracy. The higher the accuracy of the confident predictions, the smaller the number of example cells that the anatomist needs to mark to verify that accuracy.
2. **Searching for misses:** the anatomist looks for marked cells that were completely missed by the detector.
3. **Classifying the unconfident detections:** the anatomist marks all of the low confidence predictions.

Steps 1 and 2 are based on a sampling of the automatically detected cells and are therefore represent a light workload for the anatomist. Most of the work of the anatomist is in step 3. We call the ratio between the unconfident detections and the confident detections the “effective work ratio”. When the effective work ratio is small, the savings in manual work is large.

## 2 The Problem

To demonstrate our methodology, we focus on a representative challenging cell detection problem. The input consists of two 3D images of the same brain, using two florescent markers: a Neurotrace marker that functions as the counterstain, marks all neurons, and fluoresces in the blue; and a green fluorescent protein (GFP) marker that is expressed by cells of interest based on their axonal output projections [28, 31, 30]. While GFP is the main identifier of the cells of interest, the Neurotrace channel is used to eliminate false detections [32]. A typical false detection occurs when there is a GFP signal, but no indication from Neurotrace that a neuron exists in the location.

Detection of molecularly marked cells is a demanding mental process and requires discriminating cell shapes that vary greatly both within a brain and between brains. The goal is to integrate cues from the Neurotrace and GFT markers. Our estimate is that it takes a trained anatomist about 20 s to locate and mark a single cell. This involves scanning across the image of GFP marked cells and cross-checking with the co-aligned Neurotrace channel. For 10,000 marked cells this corresponds to about 50 hours. As annotating cells is tiring, anatomists typically assign no more than a few hours per day, which means that manual detection can take weeks. Our goal is to devise a methodology, consisting of both a computer algorithm and a human workflow, that exploits the fact that most marked neurons are easy to identify. Thus automation can significantly reduce the workload on the anatomist as well as the time it takes to complete annotation of a whole brain. We develop algorithms that distinguish between “sure” and “unsure” examples, automatically mark the sure detections, and identify “unsure” hard examples for further human analysis.

## 3 Methods

### 3.1 CellBoost Overview

Our system, called CellBoost, is a universal framework constructed to recognize neurons in any brain region. CellBoost consists of two main modules (Figure 2): cell segmentation module and composite detector module. Unlike traditional segmentation methods trying to solve the problem in one step, we add a recognition step as a filter for segmented candidates. Our detector aims to characterize shapes of cells and make decisions based on cell shape features. In other words, CellBoost operates similarly to people when recognizing neurons.

**Figure 2:**
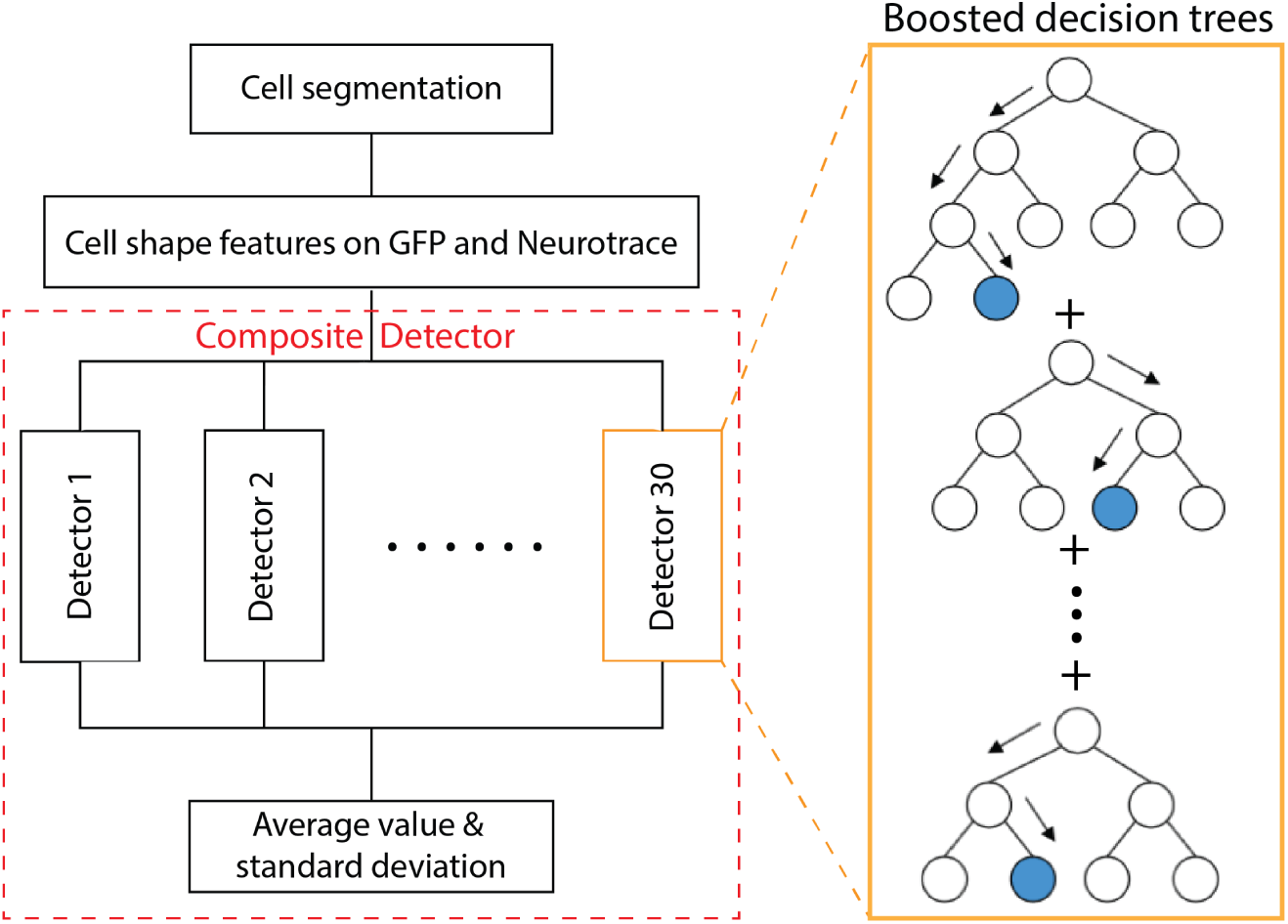
Machine Learning architecture. The system consists of four stages. The first stage is cell segmentation, which takes as input the GFP image channel and outputs a set of detection candidates. In the second stage, cell shape features are computed for each candidate. The third step consists of 30 scoring functions, each of which are boosted decision trees. The structure of a boosted decision tree is shown on the right. The fourth stage, called the composite detector, computes the mean and standard deviation of the 30 scores generated in stage 3. This mean and standard deviation are used to classify candidates into positive, negative and unsure.

**Figure 3:**
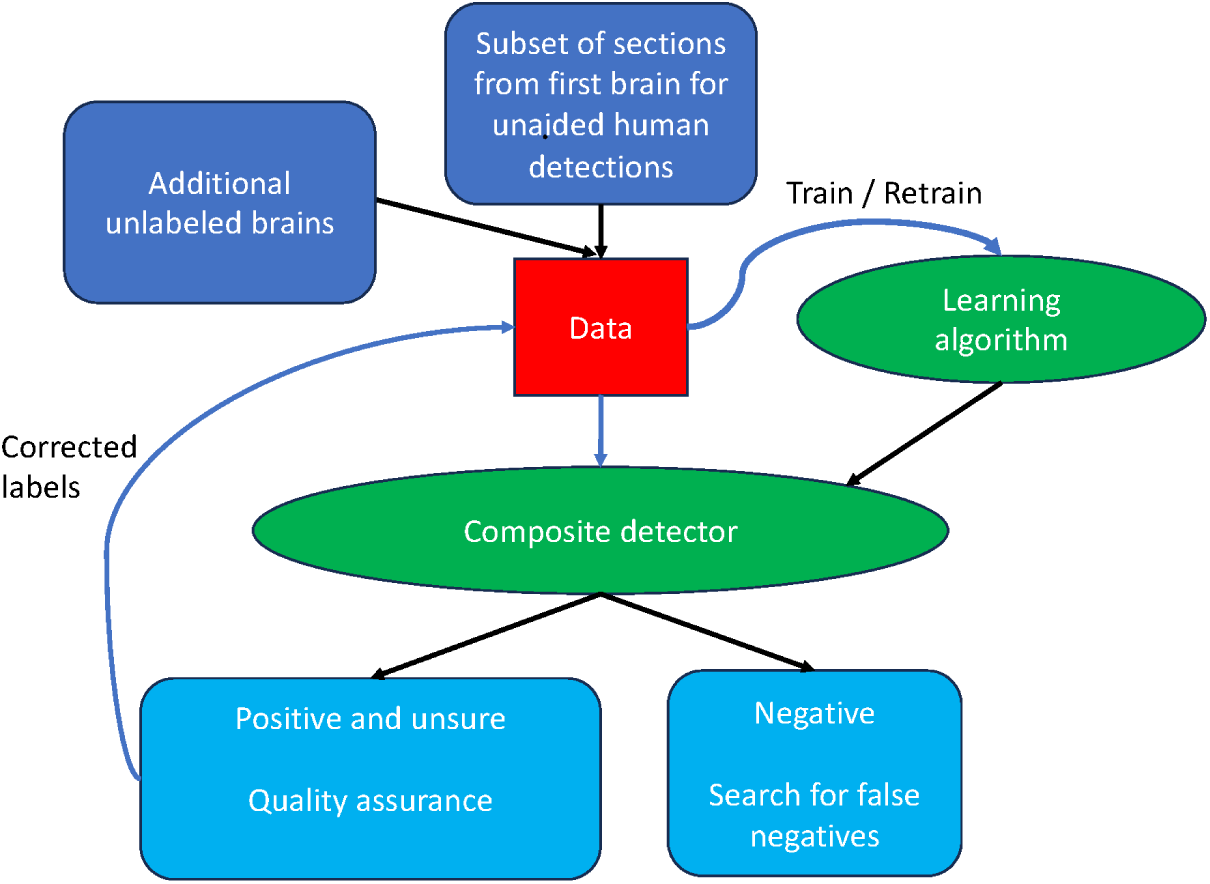
Human-machine collaboration workflow.

As a specific implementation, we apply CellBoost for the automatic recognition and annotation of labeled premotor neurons in the mouse brainstem [31]. Our detector achieves fast adaptation from few samples across different experiments.

### 3.2 Segmentation

We use adaptive threshold in the cell segmentation module, which offers the advantages of simplicity and speed. Our adaptive thresholding is conducted with the following steps on the GFP channel of the imaging data:

1. Convolve the image with a Gaussian filter. Since most of our cells are smaller than 200 pixels in length, corresponding to 65 *µ*m, we set the sigma value of the filter to be 100 and the kernel size to be 401 *×* 401 (according to the discrete Bessel approximation). This step can enhance robustness against confounds such as labeled fine neural processes.
2. Subtract the original from the convolved image.
3. Threshold the difference image with a global constant *C*. The value of *C* is set to be 2000 empirically for our data sets, whose data type is 16-bit unsigned integer.
4. Use cv2.connectedComponentsWithStats to find connected components as cell candidates in the thresholded image (Figure 1D).

### 3.3 Feature extraction

The result of the segmentation step is a list of candidates, each defined by matching regions in the GFP and Neurotrace images. Each candidate is then mapped to 40 features to characterize its shape, both in the GFP and the Neurotrace images (Table 1). These features have been handcrafted to be an over-complete representation of the candidates.

**Table 1:** Feature importance in detectors of Composite Detector G6.

| Rank | Feature name | Definition | Importance in detectors<br>(Median) |
| --- | --- | --- | --- |
| 1 | $m_{11}$ | a image moment: $\sum_x \sum_y xyI(x, y)$ | 49160.54 |
| 2 | $Xcorr\_GFP$ | mean of cross-correlation to the average<br>shape of positive cells in the GFP channel | 6735.25 |
| 3 | $m_{10}$ | a image moment: $\sum_x \sum_y xI(x, y)$ | 3352.64 |
| 4 | $m_{12}$ | a image moment: $\sum_x \sum_y xy^2I(x, y)$ | 2981.99 |
| 5 | $m_{01}$ | a image moment: $\sum_x \sum_y yI(x, y)$ | 2922.80 |
| 6 | $m_{21}$ | a image moment: $\sum_x \sum_y x^2yI(x, y)$ | 2087.11 |
| 7 | $energy\_Ntb$ | integral of squared image gradients<br>in the Neurotrace channel | 1937.21 |
| 8 | $Xcorr\_Ntb$ | mean of cross-correlation to the average<br>shape of positive cells in the Neurotrace channel | 1437.84 |
| 9 | $mu_{02}$ | a image moment: $\sum_x \sum_y (y - \bar{y})^2I(x, y)$ | 1279.08 |
| 10 | $m_{20}$ | a image moment: $\sum_x \sum_y x^2I(x, y)$ | 1249.88 |
| 11 | $energy\_GFP$ | integral of squared image gradients<br>in the GFP channel | 1147.04 |
| 12 | $h_0$ | the first Hu moment | 841.12 |
| 13 | Contrast_Ntb | $\frac{\bar{I}_{in}(Neurotrace) - \bar{I}_{all}(Neurotrace)}{\bar{I}_{in}(Neurotrace) + \bar{I}_{all}(Neurotrace)}$ | 746.79 |
| 14 | Contrast_GFP | $\frac{\bar{I}_{in}(GFP) - \bar{I}_{all}(GFP)}{\bar{I}_{in}(GFP) + \bar{I}_{all}(GFP)}$ | 731.51 |
| 15 | $h_1$ | the second Hu moment | 610.77 |
| 16 | $nu_{20}$ | a image moment: $mu_{20}/m_{00}^2$ | 355.44 |
| 17 | $mu_{11}$ | a image moment: $\sum_x \sum_y (x - \bar{x})(y - \bar{y})I(x, y)$ | 346.37 |
| 18 | height | height of a candidate | 312.76 |
| 19 | $nu_{11}$ | a image moment: $mu_{11}/m_{00}^2$ | 296.03 |
| 20 | $mu_{20}$ | a image moment: $\sum_x \sum_y (x - \bar{x})^2I(x, y)$ | 260.27 |
| 21 | $nu_{30}$ | a image moment: $mu_{30}/m_{00}^{5/2}$ | 258.57 |
| 22 | area | area of a candidate | 256.55 |
| 23 | $mu_{03}$ | a image moment: $\sum_x \sum_y (y - \bar{y})^3I(x, y)$ | 239.08 |
| 24 | $h_2$ | the third Hu moment | 232.17 |
| 25 | $mu_{21}$ | a image moment: $\sum_x \sum_y (x - \bar{x})^2(y - \bar{y})I(x, y)$ | 229.82 |
| 26 | $h_3$ | the forth Hu moment | 225.75 |
| 27 | $nu_{03}$ | a image moment: $mu_{03}/m_{00}^{5/2}$ | 225.30 |
| 28 | $h_5$ | the sixth Hu moment | 222.02 |
| 29 | $h_6$ | the seventh Hu moment | 214.95 |
| 30 | $m_{03}$ | a image moment: $\sum_x \sum_y y^3I(x, y)$ | 213.31 |
| 31 | $mu_{12}$ | a image moment: $\sum_x \sum_y (x - \bar{x})(y - \bar{y})^2I(x, y)$ | 208.33 |
| 32 | $h_4$ | the fifth Hu moment | 203.76 |
| 33 | $nu_{02}$ | a image moment: $mu_{02}/m_{00}^2$ | 202.05 |
| 34 | $mu_{30}$ | a image moment: $\sum_x \sum_y (x - \bar{x})^3I(x, y)$ | 200.54 |
| 35 | $nu_{21}$ | a image moment: $mu_{21}/m_{00}^{5/2}$ | 198.11 |
| 36 | $nu_{12}$ | a image moment: $mu_{12}/m_{00}^{5/2}$ | 188.40 |
| 37 | $m_{30}$ | a image moment: $\sum_x \sum_y x^3I(x, y)$ | 175.03 |
| 38 | width | width of a candidate | 125.28 |
| 39 | $m_{02}$ | a image moment: $\sum_x \sum_y y^2I(x, y)$ | 99.96 |
| 40 | $m_{00}$ | a image moment: $\sum_x \sum_y I(x, y)$ | 10.83 |

### 3.4 Sample

Our sample contains four brains derived from the the same experimental protocol [31], all similarly sectioned at 20 *µ*m thickness, stained, imaged at 0.325 *µ*m by 0.325 *µ*m per pixel, and aligned with Elastix [18]. Sectioning is performed with the Cryojane technique to preserves the geometry of each section [6, 22]. This approach literally fuses a section of brain tissue to a solid glass substrate and thus obviates any deformation that would add systematic noise to the data set. The somata of each cell is counterstained with Neurotrace blue and the cells retrogradely marked by the expression of GFP, as follows:

- **Brain0**: premotor jaw neurons were labeled with GFP through a retrograde transport process from the masseter muscle [31].
- **Brain1** and **Brain2**: premotor whisking neurons were labeled with GFP through a retrograde transport process from the intrinsic vibrissa muscles.
- **Brain3**: licking premotor neurons were labeled with GFP through a retrograde transport process from the genioglossus muscle in the tongue.

### 3.5 Classification and classification confidence

We obtained a total of 100,000 candidate cells through segmentation performed on every fifth section in Brain0. Out of these sections, human annotators labeled about 2,000 candidates as “positive” in an unaided mode. Subsequently, the remaining 98,000 unmarked candidates were labeled as “negative” examples. These 100,000 examples formed the intial training set. Each example is then described by a vector of the 40 features (Table 1). We use a composite detector of 30 individual detectors (Figure 2). Each detector is a boosted tree [11, 5] generated using the same training data but with a different random seed. The outputs of the individual detectors are combined using bagging-style averaging [7]. Two quantities are computed based on the composite detector. One quantity is the mean of the scores and the other is the standard deviation of the scores. Both are indicative of the prediction confidence. The mean is indicative of the prediction margin [1], while the standard deviation is inversely proportional to the stability across random seeds [2]. We use the mean to partition “sure” from “unsure” examples and the standard deviation to verify that candidates with low mean scores are unstable. To compute the classification of a candidate into “positive” and “negative” “sure” detections and “unsure” detections we use two thresholds. These thresholds are chosen based on the scatter plots of mean and standard deviation (Figure 4).

**Figure 4:**
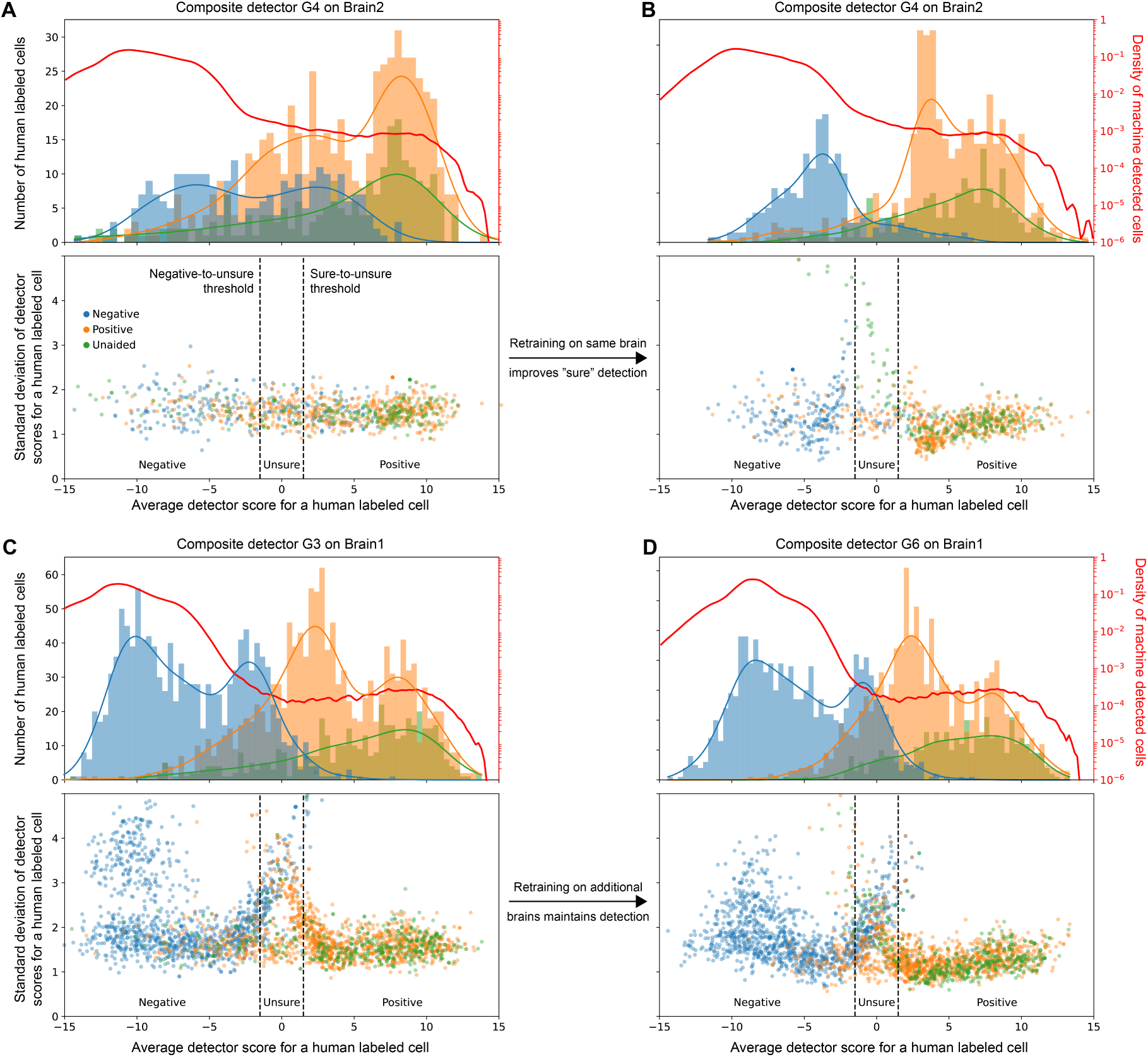
Performance of composite detector: Each panel in this figure summarizes the performance of a single composite detector. The horizontal axis corresponds to the average score and is divided into three regions: Positive, Unsure, and Negative. The vertical axis of the scatter plot corresponds to the standard deviation of detector scores. The dots correspond to human labeled examples which labeled by three colors: blue and orange dots corresponds to cells that have been labeled positive and negative in QC, while green dots correspond to cells that have been detected by a human unaided. In the histogram plot, we show the quantity distribution of these human labeled examples. Besides, the bold red curve and the corresponding right axis describe the density of the scores of the candidates. Note there are two orders of magnitude difference between the density of the negative and the positive candidates. **(A,B)** Here we observe the improvement in performance on Brain2 before and after training on Brain2. **(C,D)** Here we observe the stability in performance on Brain1 maintained after training on brains other than Brain1.

### 3.6 Human machine cooperation

The human annotation effort was most extensive with the first of the four brains, with much smaller efforts required for the other brains. Some of the sections from the first brain are annotated by a human “unaided”, which means that the human has to annotate without the aid of the computer. Unaided annotation is the most labor intensive step and requires about 6 to 10 minutes per brain section.

After the composite detector, trained on the initial training set, is applied to all sections of the brain, the output is checked by humans for QC. This analysis is significantly less labor intensive that unaided annotation and proceeds as:

- Equal size samples are selected from the “positive” and “unsure” subsets and mixed. The candidates labeled “unsure” are too hard for the machine to call and are left for humans to label. A human then performs annotation to verify that the accuracy of the “positive” is high. This annotation is more efficient than the unaided because the computer centers the display at the candidate cell, and the human has to only answer yes or no.
- The “sure” detections, together with the manual corrections, are fed back to the training data and used to retrain the detector.
- The “negative” subset is typically much larger than the rest. To ensure that false “negatives” are rare, the human performs a search where s/he is looking for undetected cells.
- The ratio between the sizes of the sure and unsure sets, as noted, is the “work saving ratio”. Human marking is reduced to the following two tasks by our system:
- **Quality control:** Marking a sample of the 250 “positive” and 250 “unsure” to verify the accuracy of the “positive” and “unsure”.
- **Unaided annotation:** Marking five sections in an “unaided” annotation mode to estimate the false “negatives” of the detector.

After completing these tasks, the user has a good estimate of the “false positive” and “false negative” rates. If these are sufficiently low, the task is done and the “positive” detections are accepted for the neuroanatomical analysis. If the QC performance of the composite detector on a brain is insufficient, then the composite detector is retrained by using the QC examples together with examples that are high confidence “positive” and high confidence “negative”.

## 4 Results

### 4.1 Segmentation

Our relatively simple segmentation method manages to catch nearly all the marked cells. In general, when parameters are set to ensure low numbers of false negatives, the number of false positives is similar across many segmentation methods. Specifically, our experiments show that the adaptive threshold we use for CellBoost performs similarly to established methods such as Cellpose [29] (Figure 1), a popular cell segmentation method based on deep learning methods with NNs. Notably, Cellpose requires annotating cell boundaries to retrain models, a process that demands extensive manual labor.

In practice, Cellpose overlooks some true positive cases when two or three cells overlap in space or are closely situated (yellow rectangle in Figure 1). Our framework uses a simple and efficient segmentation step followed by a machine learning classification step to detect the true positives.

### 4.2 Machine Learning Performance

Our composite detector module consists of an ensemble of boosted decision tree models [5] and classifies cell candidates as “positive” or “negative” for confident detections, and “unsure” for unconfident detections. We trained and tested six generations of composite detectors. The initial composite detector begins its training with **Brain0**. Here a human anatomist performed unaided annotation of every fourth brain section to mark the GFP expressing premotor neurons. Subsequently, the detector underwent testing and retraining on **Brain1**, **Brain2**, and **Brain3**, in that order.

#### 4.2.1 Detector Accuracy

We conducted two tasks, i.e., quality control (QC) and unaided annotation. These tasks require a small sample of manual annotation and are efficient to conduct. The quantitative results are:

- **Quality Control:** It took an average of 108 minutes to complete QC of 500 samples, corresponding to 13 seconds per cell. The error rates of “positive” detections are 4.4%, 12.8% and 23.2%, respectively. The human disagreement rates of “unsure” detections are 9.2%, 16.8%, and 24.4%, respectively. The error rates of “positive” detection appear correlated to the human disagreement rates of “unsure” detections.
- **Unaided Annotation:** It took an average of 35 minutes to assess five sections with labeled cells, corresponding to 5.4 seconds per cell. The “false negative” rates are 7.8%, 8.3%, 9.0% respectively. The “false positive” rates are 0.03% (12/40448), 0.70% (43/6550), 0.14% (92/66242), respectively. Note that they are all rates are close to 0% because of the large number of “negative” detections.

#### 4.2.2 Improvement by Retraining

We provide scatter plots based on the mean and standard deviation of detection scores in our composite detector to visualize the improvement by retraining. The performance of the classification stage is summarized in the plots of Figure 4 and compared with human performance. We make the following observations:

- Most of the segmented candidates receive negative scores. This occurs since most cell candidates correspond to small regions that are not cells.
- The QC labeling shows a low error rate on the confident “positive” and the confident “negative” classifications.
- The unaided labeling indicates that most of the unaided labels are classified as “positive”. A small fraction are “unsure” and about 5% are omitted.
- The relationship between the average and standard deviation of the detector scores shows that, in general, the margin and the standard deviation are closely related. But when many examples are annotated, as in Brain1, there are many unstable “negative” examples that are still recognized as confidently “negative”.

We make the following observations regarding retraining based on our training sets:

- **Brain1:** This brain was analyzed using an early detector (G3) and the performance was compared to that of a later detector (G6) that was trained on additional brains. In this case, we observe that the performance on Brain1 did not degrade by adding training data from additional brains. This gives evidence that retraining a detector on additional brains does not degrade the performance on prior brains.
- **Brain2:** This brain was analyzed using detector data from other brains resulting in a well trained detector (G4). To improve the performance, a new detector (G5) was trained on the existing data set with the addition of 500 cells from the QC of Brain2. Comparing panels A and B in Figure 4, we see that a significant improvement in performance was achieved using a small number of training examples.

It is not surprising that this improves performance on additional brains. More significant is that the performance on initially analyzed brain shows an improvement as well.

### 4.3 Feature selection

Our method designed a group of features characterizing cell shapes and provided feedback regarding the importance of these features utilized in cell detection, which is not available in NNs. The relative importance of the features that are used in the composite detectors is depicted in Figure 5. The importance is defined as the total gain across all splits the feature is used in a decision tree model. The main observation is that the the most important features are low-order moments of the GFP image channel and the cross-correlation coefficient (defined in Table 1) calculated from the GFP channel as well as from the Neurotrace channel. As anticipated, features from the Neurotrace channel play a pivotal role. Unexpectedly, central moments and other invariant moments appear to be less important than raw moments. This indicates that scale and rotation might significantly influence the detection process.

**Figure 5:**
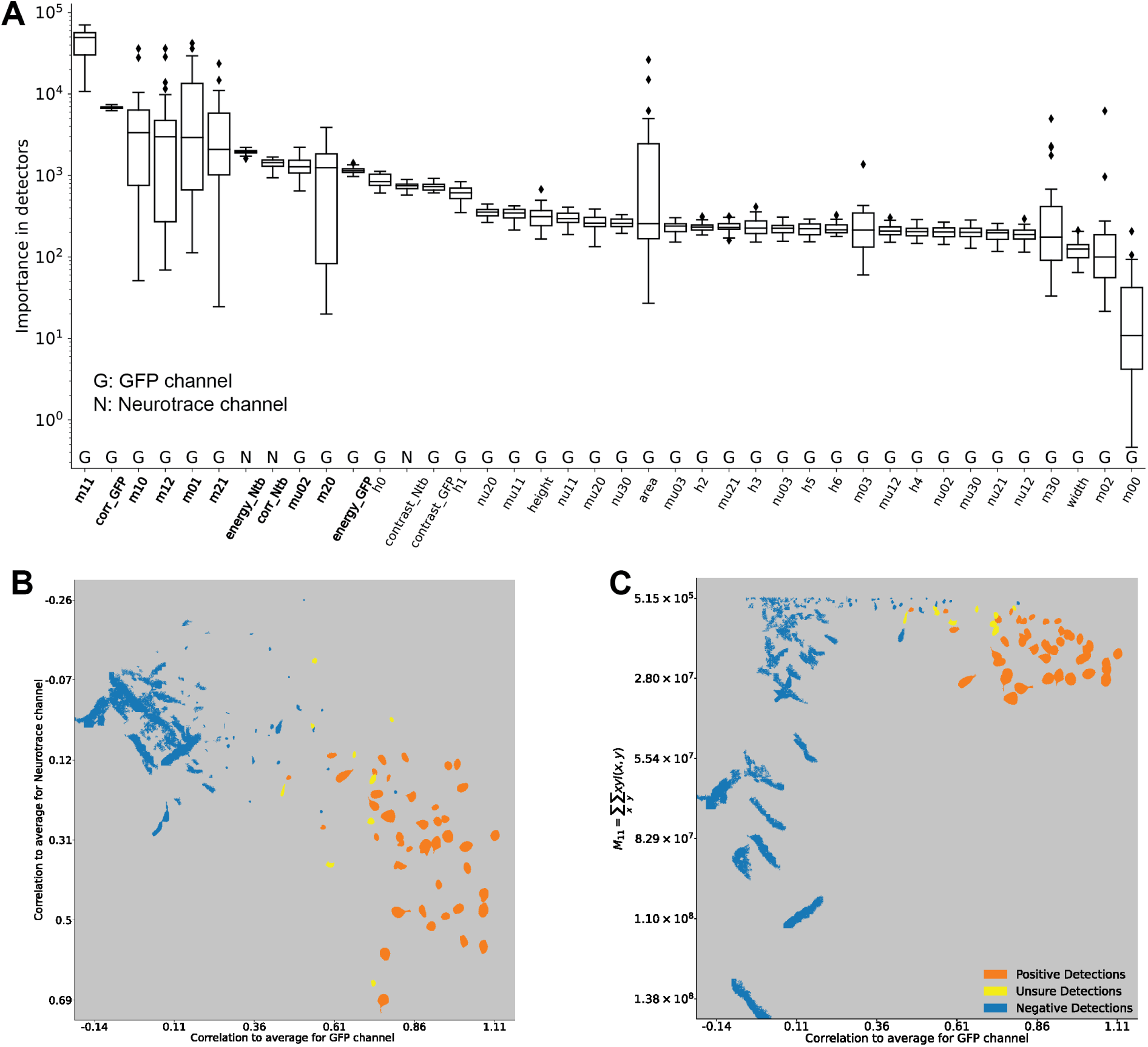
The important shape features. The boosting algorithm uses as input all 40 features described in Table 1. However, some of the features carry more weight than others. **(A)** Display of feature importance, in decreasing order of weight. A box plot is used to show the distribution of the weight across the 30 bagged copies of the detector that are incorporated into the combined detector. **(B,C)** Scatter plots of the real shapes of the GFP images of the cells distributed according to two high-weight features. In (B) the horizontal distribution is according to the across-correlation to the average shape in the GFP channel, while the vertical is the same for the Neurotrace channel. In (C) the horizontal is the same as (B) while the vertical corresponds to moment 11 (*m*_11_).

Detailed scatter plots for a sample of cell shapes comparing GFP cross-correlation coefficients to Neurotrace cross-correlation coefficients (Figure 5B) and GFP cross-correlation coefficient to second-order moment *m*_11_ (Figure 5C). We observe distinct clusters corresponding to “positive” and “negative” detections, with a few “unsure” detections situated between the two clusters. Cell shapes similar to the average shape of premotor neurons are associated with high GFP cross-correlation coefficients. This feature exhibits an outstanding capability to distinguish between “positive” and “negative” detections. The clustering observed in the scatter plots justifies the large weights of the corresponding features in Figure 5A.

### 4.4 Comparative performance

We found the consistence between our automated detections and human annotations by checking characteristics of samples in the QC process (Figure 6). Most fluorescently marked premotor neurons are situated in the brainstem. Figure 6A identifies multiple regions in a brainstem section with clear and robust fluorescent signals; two prominant regions are the spinal trigeninal subnuclei oralis (SpVO) and interpolaris (SpVI) and the Kolliker-Fuse nucleus (KF), and, as expected, no detection in the vibrissa facial motonucleus (vFN) [31]. Correspondingly, Figure 6B shows that our “positive” detections are primarily localized in these two regions. Conversely, “negative” detections are observed throughout the entire brain, consistent with the fact that most cell candidates are not real cells. An area is magnified to show typical premotor neurons, characterized by apparent GFP-expressing shapes accompanied by distinct black cell bodies. As shown in Figure 6C, our detector identifies these neurons and categorizes other minor stained objects as negative. Three of these detections were selected as samples for QC and there was a consensus among our human annotators regarding the detection outcomes.

**Figure 6:**
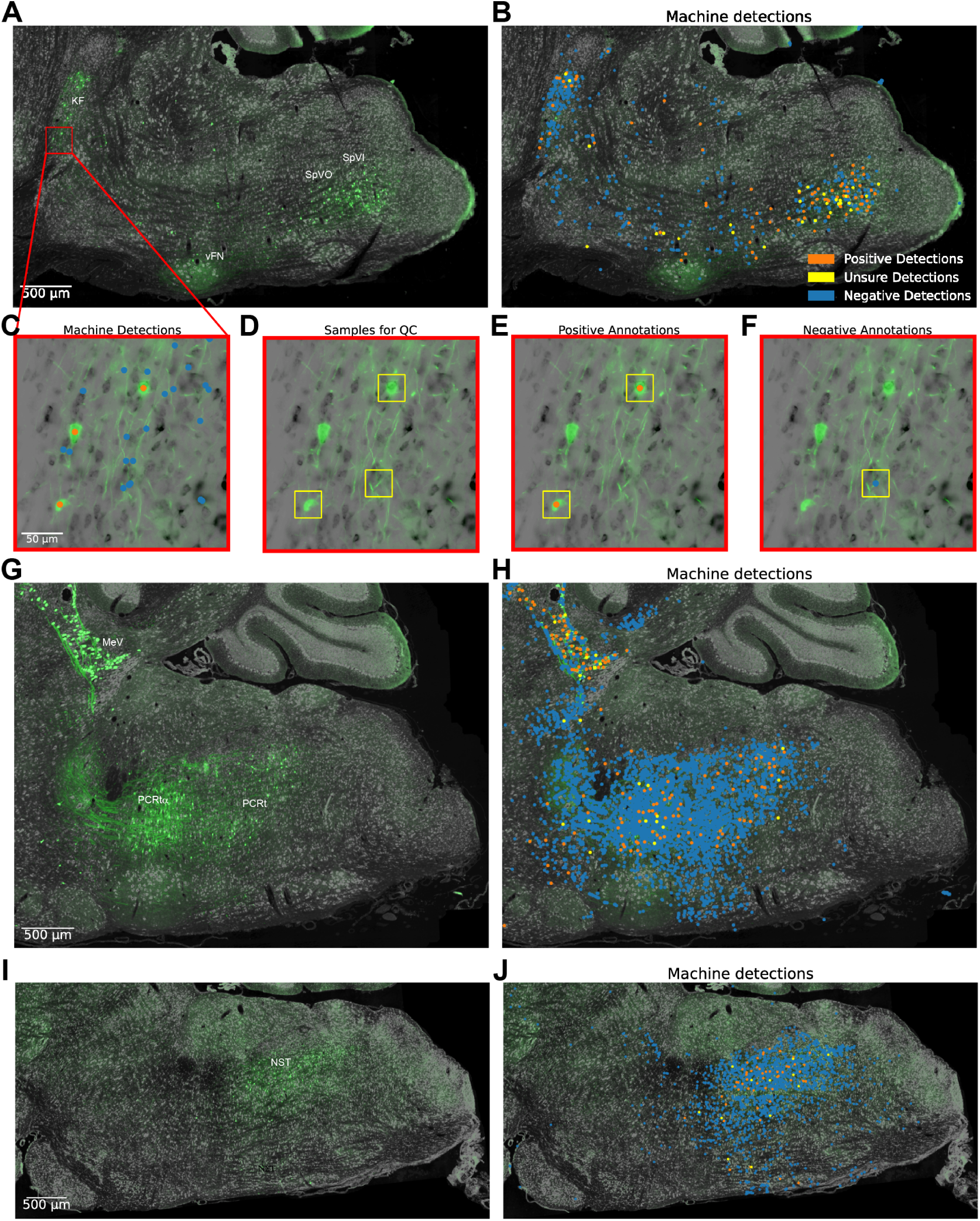
Overview of the performance. **(A)** A section of the brainstem stained with GFP and Neurotrace from Brain1, where intrinsic vibrissa muscles were injected. Abbreviations: SpVI = spinal trigeminal subnucleus interpolaris; SpVO = spinal trigeminal subnucleus oralis; KF = Kolliker-Fuse; vFN = vibrissa subregion of facial motor nucleus. **(B)** Detections classified as “positive”, “unsure”, and “negative”. This illustrates the spatial distribution of machine detections. **(C)** A magnified area highlighting machine detections of all confidence levels. **(D-F)** Randomly selected samples from detections for QC, with positive (E) and negative (F) annotations provided for clarity. **(G-H)** A section of the brainstem stained with GFP and Neurotrace from Brain0, where the masseter muscle was injected. Abbreviations: PCRt = parvocellular region of the reticular formation; PCRt*α* = *α* regionof PCRt. **(I-J)** A section of the brainstem stained with GFP and Neurotrace from Brain3, where the genioglossis muscle was injected. Abbreviations: NST = nucleus of the solitary tract.

The disagreements between two human annotators is highlighted in Figure 7. For one QC sample, an annotation could either be “positive” or “negative”, yielding four potential outcomes between two human annotators. We separate GFP and Neurotrace image channels to explore characteristics of these outcomes.

- A “sure” “positive” example labeled “positive” by both annotators (Figure 7A). This is a typical premotor neuron, evidenced by a green-stained shape with clear boundaries and a distinct gray cell body. Notably, our detector awards it a confident positive score.
- A “sure” “negative” example labeled “negative” by both annotators (Figure 7B). Despite the presence of a green-stained shape in the GFP channel, the corresponding location in the Neurotrace channel offers no indication that a neuron exists. In response, our detector assigns this cell candidate a confident “negative” score.
- Two hard cases, with different annotations from two human annotators (Figure 7C, D). Both cell candidates exhibit green-stained shapes with blurry boundaries, while vague signals can be found in the Neurotrace channel. These traits lead to disagreement between annotators and also an “unsure” detection by our detector.

**Figure 7:**
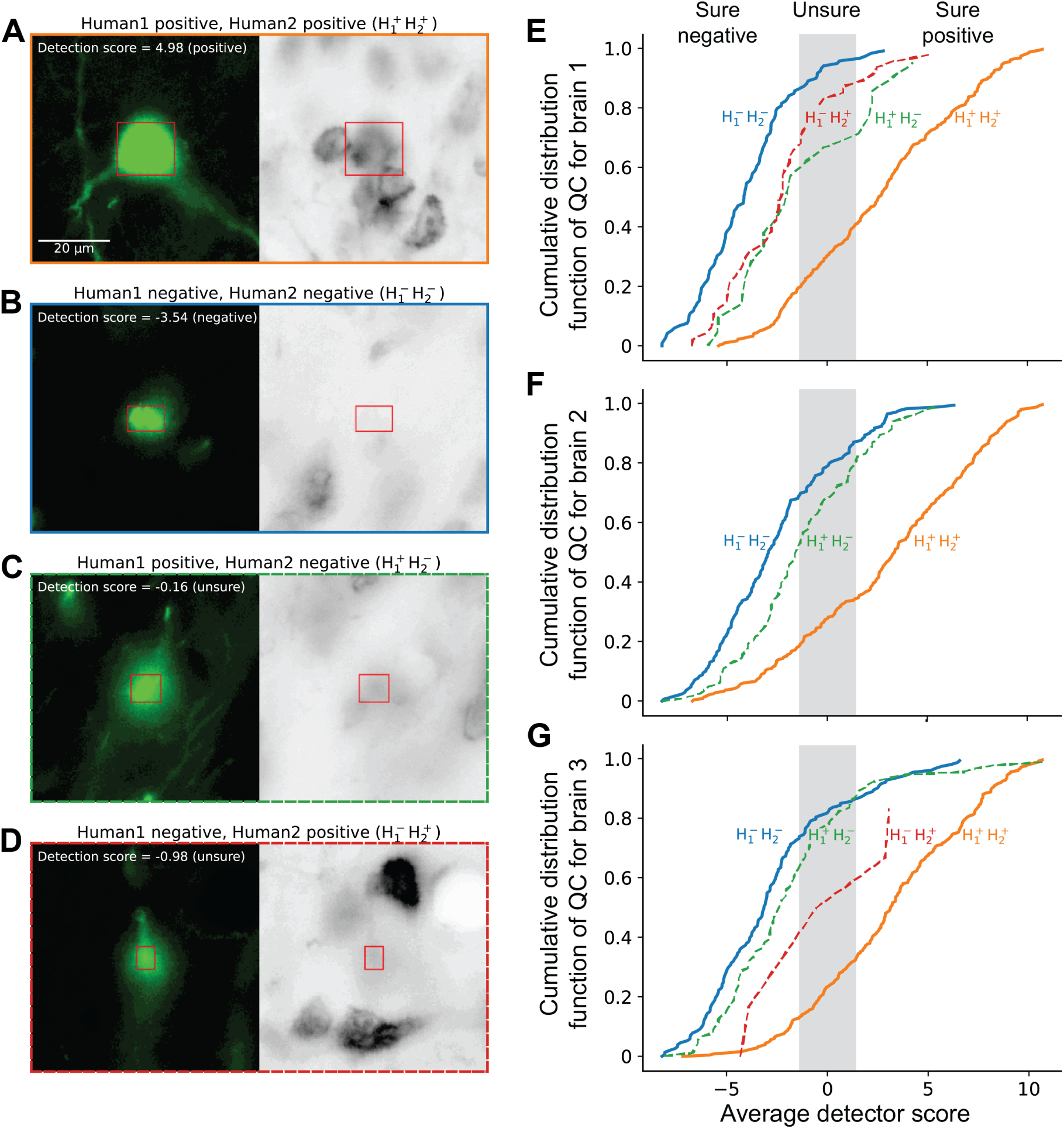
Human performance. **(A,B,C,D)** Depiction of the four possible outcomes between two human annotators, with the GFP and Neurotrace image channels displayed separately to underscore their roles in recognition. **(E,F,G)** Performance summary of the two human annotators across QC tests for three distinct brains, with CDFs plotted for the four potential outcomes based on average detector scores. For brain 1, the case distribution is as follows: 313 *H*_1_^+^*H*_2_^+^, 119 *H*_1_*^−^H*_2_*^−^*, 21 *H*_1_^+^*H*_2_*^−^* and 47 *H*_1_*^−^H*_2_^+^ respectively. For brain 2, there are 245 *H*_1_^+^*H*_2_^+^, 169 *H*_1_*^−^H*_2_*^−^*, 83 *H*_1_^+^*H*_2_*^−^* and 3 *H*_1_*^−^H*_2_^+^ respectively. For brain 3, there are 250 *H*_1_^+^*H*_2_^+^, 153 *H*_1_*^−^H*_2_*^−^*, 91 *H*_1_^+^*H*_2_*^−^* and 6 *H*_1_*^−^H*_2_^+^ respectively. Notably, there is an approximate 15% to 20% disagreement rate between human annotators. The disagreements are largely within the unsure region.

Neurotrace Nissl staining is a very useful histological method to facilitate the identification of cell populations. Our anatomists predominantly utilize the Neurotrace channel to verify the existence of premotor neurons. Therefore, vague signals in this channel can pose challenges in decision-making. If annotators establish individualized criteria, their discrepancies might stem from systematic errors. We observe that nearly 15% to 20% of QC samples for each brain display divergent decisions from the two annotators (Figure 7E-G). Strikingly, of the total disagreement cases, 83 out of 86 (Figure 7F) and 91 out of 97 (Figure 7G) come from the same category, labeled “positive” by one human annotator while “negative” by the other. This indicates that some disagreements may be attributed to a systematic bias, with image quality playing a pivotal role in the annotators’ judgment.

We checked if examples identified as hard by the algorithm are also hard for human annotators. We assigned two human annotators to mark a sample of hard examples and counted the disagreements. Our results show a human disagreement rate of about 20%. The human disagreement rates of “unsure” detections are 12.24%, 22.67%, and 32.21%, respectively, in this test.

The level of agreement between human annotators is studied in inter-rater experiments [12]. Inter-rater agreement measures the rate of agreement between marked cells chosen by different annotators. A common way to quantify the agreement between two raters is the Cohen’s *κ* (kappa) coefficient [19].^2^ The human disagreement rate of about 20% in our experiment corresponds to *κ* = 0.8, on the border of what is considered perfect agreement and substantial agreement.

## 5 Discussion

A challenge for current methodology in anatomical studies is the identification of brain neurons. Classically, this relies on labor intensive processes. The automation of cell detection has focused on techniques to completely replace the human annotator, even for data sets with ambiguously stained cells. Yet this can lead to false detection. The central motivation of our work is to develop tools to leverage human expertise rather than attempt to replace it. This provides a means to minimize false detection by focusing the human effort on the small number of ambiguous cases.

The first novelty of our pipeline is that it partitions the candidate labeled cells into two classes based on reliability. One is a “sure” or reliable categorization as “positve” or “negative” based on the assessment of molecular marking. The second is an “unsure” or borderline category reflecting ambiguity of the molecular marking. The utility of this approach is that examples that are difficult to classify then to end up in the “unsure” category. This results in increased accuracy of the “positive” and “negative” classes. Implementing this scheme requires a rigorous metric of confidence that is applicable to our algorithm [2, 1].

Our study reveals that human annotators disagree with each other on 10 - 20 percent of their annotations. Moreover, these disagreements are concentrated in the candidates that receive an “unsure” label by the algorithm. This means that the set of examples for which the algorithm outputs “unsure” has a significant overlap with the set of examples on which there is disagreement between human annotators. The cells in the “unsure” set are atypically shaped cells or a result of systematic biases such as artifacts in the fluorescent channels.

A second novelty of our pipeline is the reduction of manual labor. We use two annotation protocols. In the first protocol called “unaided”, the human views a set of molecularly marked sections and is asked to find all of the marked cells in those sections. The unaided task is the baseline that is used when no composite detector has yet been trained. In the second protocol, called “QC”, the human is presented with the location of candidate cells and is asked to label them “positive” or “negative”.

The processing of the brain consists of two stages, a training stage and a maintenance stage. In the training stage, the first brain is annotated using the unaided protocol. This is a significant task that can take several weeks. The detections from the unaided protocol are transformed into several thousands positive examples, to which a sample of undetected candidates are added as negative examples. This training set is used to train and test the composite detector. In the maintenance stage the composite detector is applied to the brains sequentially to split the candidates into “positive”, “negative” and “unsure”. The main human task at this stage brains is to perform QC to verify that the “positives” are correct. This is done using a sample of 500 candidates that are typically labeled by a human in two hours. The outcome of the QC is either that the accuracy is sufficient or insufficient. For the former case the “positive” labeled cells are output and added to the training set. In the latter case the outcome of the QC is added to the training set to generate a more accurate composite detector. Our experiments show that one to two QC cycles, hours versus weeks of work per brain, are sufficient to achieve a high accuracy of labeling molecularly marked cells.

The system we describe can be extended to count marked neurons within specific anatomically and/or functionally distinct regions. This requires the addition of three-dimensional boundaries of the regions, as may be implemented by systematically mapping the anatomical data to a known atlas [35, 6, 10, 14, 16]. Here our focus was on the application of the Boosting method to cell identification, for which we used four brains with retrogradely labeled premotor neurons starting from individual muscles involved in orofacial active sensation [31, 15]. This led to a tool that leveraged human expertise rather than attempted to replace them.

Our pipeline can be applied to other stain combinations of molecular markers, brain regions, and species. The benefit of using our pipeline is that only one brain has to be labeled “unaided”, requiring weeks of manual work. Other brains of the same type require only a few cycles of QC, each of which can be completed in two hours. We hope that other neuroscience laboratories would find our pipeline useful with the increased popularity of high-throughput microscopy.

## 6 Code availability

This software is fully open-source, along with our custom visualization and annotation tools built upon Neuroglancer, a WebGL-based software.

Cell detector: https://github.com/ActiveBrainAtlas2/cell extractor Neuroglancer: https://github.com/ActiveBrainAtlas2/neuroglancer

## Acknowledgements

We thank Harvey Karten and Samuel Wang for useful discussions, Julian Dukes and Marissa Moreno for their role as annotators, Hannah Liechty for assistance in scanning, and Sincheng Huang for comments on an early version of the manuscript. This work was supported by the National Institutes of Health (grants U19 NS107466 and RF1 MH128776).

## A Methods details

### A.1 Boosting and sparse representations

An important part of the design of any learning algorithm is finding a representation of input feature vectors that captures the aspects that are most relevant for the classification task. In some situations deep Neural Networks can find internal representations autonomously, without human intervention. However, a close look at the design of alpha-Go [26] reveals the high level of human expertise was used to design the features used by the Neural Network.

Boosting [11, 23] is another popular learning algorithm which combines a large number of so-called “weak” rules to construct a single “strong” rule. Here we follow an approach to feature detection for boosting that can be described as the kitchen sink approach. This approach starts by the human constructing a very large number of candidate rules. The boosting algorithm performs both feature selection i.e. finding the rules that provide significant information about the labeled cell, as well as feature weighting and combination, i.e. finding how to combine the informative features to predict the labeled cell.^3^

In general, increasing the number of features or rules increases the danger of over-fitting. However, as shown in reference [1], the number of features has only a small influence on over-fitting. Rather, it was shown that the distribution of the large normalized margin guarantees low over-fitting even if the number of features goes to infinity.

## Footnotes

1 ^1^We restrict ourselves to binary labels. We use the term “example” to the pair (”instance”,”label”)

2 *κ* is computed from two more basic quantities: 0 *≤ a ≤* 1 is the fraction of cases on which the two raters agree, and 0 *≤ c ≤* 1 is the fraction of agreements that would occur by chance if the two annotators are statistically independent. The definition of kappa is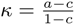. If *κ* = 1, the anatomists always agree; if *κ* = 0, the rate of agreement corresponds to chance; and if *κ <* 0, then the rate of agreement is lower than chance, i.e. the two anatomists tend to have different opinions. An interpretation of *κ* recommended by Cohen [19] is: *κ ≤* 0: no agreement, 0 *< κ ≤* 0.20: none to slight agreement, 0.2 *< κ ≤* 0.40: fair agreement, 0.4 *< κ ≤* 0.60: moderate agreement, 0.6 *< κ ≤* 0.80: substantial agreement, and 0.8 *< κ ≤* 1.00: perfect agreement.

3 Popular boosting software, such as XGBoost and LightGBM, use decision trees to combine the features.

